# Gene regulatory networks and the organization of cell differentiation in multicellular organisms

**DOI:** 10.64898/2026.11.18.767571

**Authors:** L. Fernando Meraz-Segura, A. Chárbel Guerrero-Vaca, Rafael Alcaraz Barrera, Carlos Espinosa-Soto

## Abstract

Across their development, clonal multicellular organisms generate new cell types from less differentiated cell types. Differentiation pathways arrange into a hierarchical, tree-like organization: regressions from specialized to undifferentiated cell types rarely occur and undifferentiated cell types often have the potential to produce, when subject to distinct stimuli, two or more differently specialized cell types. How cell type transitions acquire such a peculiar disposition requires an explanation that must reside in the properties of gene regulatory networks (GRNs); after all, GRNs orchestrate the changes in gene activity that define cell types. We studied vast numbers of small dynamic model networks to search for elementary GRNs that could yield the simplest hierarchical, developmental-like, differentiation patterns. We found that only a small minority of GRNs are compatible with hierarchical cell type transitions. Importantly, GRNs that yielded developmental-like differentiation patterns shared common traits: One is the presence of tangles of cycles of positive interactions. Genes in such a tangle produce an all-or-none one-way response to a signal. Another such trait is mutual inhibition between genes associated to different specialized cell types, enforcing their mutually exclusive activation. Moreover, these regulatory patterns are more abundant in published experimentally sustained developmental GRNs than as expected by chance. In addition, we show that it is easy to maintain hierarchical cellular differentiation throughout evolution, even when undergoing changes in gene regulatory interactions and developmental mechanisms. Our work supports the prevalence of design principles that organize hierarchically differentiation pathways in clonal multicellular organisms.

## 1 Introduction

Cell differentiation is a central process in development; it implies changes in a cell’s properties and the acquisition of a new cell identity (Wolpert et al., 2019). These transformations generate all the distinct cell types in clonal multicellular organisms like plants and animals (Brunet and King, 2017).

Cell differentiation is usually described as a hierarchical process (Garcia-Bellido and Ripoll, 1978; Kauffman, 1982; Stent, 1985; Zhou and Huang, 2011).The idea of hierarchy implies several intertwined properties (Rasskin Gutman and Esteve Altava, 2009; Corominas-Murtra et al., 2011, 2013; Cheng et al., 2015; Zhang et al., 2017); here we focus on two of them: progressivity and divergence. By progressivity in cell differentiation we allude to directional cell transitions. Typically, undifferentiated cell types have the potential to produce, under appropriate conditions, more specialized cell types under appropriate conditions (Sulston et al., 1983; Plass et al., 2018; Wolpert et al., 2019). Were regressions from specialized to undifferentiated cell types frequent, the map of all possible cell type transitions would contain many circular paths, say from cell type *A* to *B*, then to *C* and back to *A*. However, these kinds of cycles do not usually appear in regular development (Kondo et al., 2003; Clevers et al., 2017).

Cell differentiation is divergent because undifferentiated cell types often have the potential to produce, when subject to distinct stimuli, multiple differently specialized cell types (Garcia-Bellido and Ripoll, 1978; Sulston et al., 1983; Gordon and Gordon, 2016; Laurenti and Göttgens, 2018; Soldatov et al., 2019; Siebert et al., 2019; Mittnenzweig et al., 2021) Thus, the number of different cell types in an organism increases as its development proceeds. Convergence, in which the same cell type can arise from two distinct cell types, can also occur (Graham, 2010), but is far less frequent than divergence. Progressivity and divergence in cellular differentiation reflect a high degree of order in organismal development (Kauffman, 1982; Stent, 1985). How do organisms achieve such an idiosincratic structure for their cell lineages?

In multicellular organisms, an undifferentiated cell may receive a signal that somehow perturbs its cellular state (Xiong and Ferrell Jr, 2003; Engler et al., 2006; Nemashkalo et al., 2017; Wolpert et al., 2019; Mayor, 2019; Barresi and Gilbert, 2023). Cell differentiation requires such a signal to activate particular genetic programs through the expression of specific genes and repression of others (Vierbuchen and Wernig, 2011; Davis and Rebay, 2017; Mayor, 2019; Wolpert et al., 2019). Gene regulatory networks (GRNs) coordinate changes in gene activity; GRNs are sets of genes whose activity regulate the activity of other genes in the same set (Peter and Davidson, 2015). Regulatory interactions among genes define the conditions that alter a gene’s activity. The arrangement and properties of the network’s regulatory interactions define which genes will eventually remain active and which will eventually turn off in response to a signal. GRNs not only define the different stable gene activity patterns that a cell can adopt; they also determine which cell type transition follows after perturbation of a given cell type (Enver et al., 2009; Bornholdt and Kauffman, 2019). Therefore, we contend that explaining progressivity and divergence in cell lineages of multicellular organisms requires understanding how GRN’s properties affect cell type transitions.

Research efforts spread across more than half a century have already accomplished significant advances. We now have it clear that positive feedback loops, in which a gene’s activity promotes itself directly or indirectly, is a requisite for the ability to yield multiple stable gene activity patterns (Thomas, 1991). Indeed, such feedback loops can play an important role in network stability (Pinho et al., 2014) and in developmental lock-in (Davidson et al., 2002; Xiong and Ferrell Jr, 2003); for example, transient activation of some genes in a positive feedback loop can result in sustained activity of all the genes in the loop (Xiong and Ferrell Jr, 2003).

Other small interaction patterns also seem to be common in GRNs involved in cellular differentiation. Reviewing transdifferentiation experiments, Graf and Enver noted that a gene regulatory interaction pattern called *self-activating toggle switch* appeared frequently among genes involved in differentiation (Graf and Enver, 2009). This interaction pattern consists of two mutually repressing genes that self-activate (Huang et al., 2007; Chickarmane and Peterson, 2008; Jia et al., 2017; Duddu et al., 2026). Its structure implies a choice between two alternative states in which either of the self-activating genes is active. Hence, it might play an important role in developmental choices (Graf and Enver, 2009). In fact, research has provided further support to the abundance of this interaction pattern in development (e.g. Chickarmane and Peterson, 2008; Bessonnard et al., 2014; Zhou et al., 2016) and their ability to yield bifurcating differentiation patterns (Huang et al., 2007; Chickarmane and Peterson, 2008; Wang et al., 2010; Bhattacharya et al., 2011; Bessonnard et al., 2014; Zhou et al., 2016; Jia et al., 2017; Duddu et al., 2026).

We may consider a generalized version of the self-activating toggle switch: sets of genes that maintain their joint expression through cross-activation and that uphold reciprocal inhibitory interactions with other such sets of genes (Huang et al., 2007; Shyam et al., 2025). Such an arrangement of gene regulatory interactions also seems to be abundant in developmental GRNs (Hari et al., 2022; Shyam et al., 2025). Moreover, sets of cooperatively cross-activating genes that repress the activity of other such sets may correspond to *character identity networks* that confer cell type identities and that set cell types as evolutionary units (Wagner, 2014; Pavlčev and Widder, 2015; Arendt et al., 2016).

Notwithstanding the many significant advances on the subject, there are many open questions on how gene regulatory networks determine progressivity and divergence in cell differentiation: What are the simplest GRNs that can yield progressive and divergent differentiation patterns? Do all of them include positive feedback and mutual repression or are there multiple different ways to attain progressivity and divergence in cell differentiation? Does nearly any gene regulatory network can produce hierarchy in cell differentiation, or is hierarchical differentiation an attribute associated to only a few of all possible gene regulatory networks? What properties in gene regulatory networks define hierarchy in cell differentiation? To start sketching some answers to these questions, we studied a version of a model that has been very useful to address many problems in evolutionary and developmental biology (e.g. Wagner, 1994, 1996; Espinosa-Soto, 2025). As usual when modeling the role of gene activity in cell differentiation (Kauffman, 1969; Espinosa-Soto et al., 2004; Enver et al., 2009; Bornholdt and Kauffman, 2019), we identified cell types with stable gene activity patterns. Thus, we we were able to follow a network’s dynamics of gene expression to distinguish what cell type transitions were compatible with a gene regulatory network after small gene activity perturbations. We studied all possible gene regulatory networks containing two, three or four genes and samples of larger networks to find out which of them yielded cell type transitions displaying progressivity and divergence. We found that the vast majority of gene regulatory networks are not compatible with hierarchical cell type transitions. However, we were able to identify small networks that produced progressive or progressive and divergent transition maps and their common traits: one of these traits is the arrangement of genes in entangled cycles of positive interactions that yield all-or-none non-reversible responses. Another one is the reciprocal repression between genes whose expression marks different specialized cell types, resulting in their mutually exclusive upregulation. Larger GRNs that produce the same hierarchical differentiation patterns follow the same rules. We also found that it is easy to maintain a hierarchical cellular differentiation through evolution, even when undergoing changes in gene regulatory interactions and developmental mechanisms. In addition, we found that the same regulatory patterns are more frequent than expected by chance in published developmental GRNs that play a role in cellular differentiation in a diverse group of plants and animals. In sum, our work substantiates the existence of design principles that clonal multicellular organisms utilize to organize hierarchically their differentiation pathways.

## Results

### From gene regulatory networks to cell type transition maps

The main goal of this work is to evaluate how genetically determined regulatory interactions constrain developmental possibilities in cell differentiation. We build simple models of gene regulatory networks that drive the dynamics of gene activity, considering the simplifying assumption that a cell’s properties are entirely defined by gene activity patterns. As often in models for development, we associate a network’s fixed-point attractors, its stable gene activity patterns, with an organism’s different discrete cell types (Kauffman, 1991; Kauffman et al., 2003; Wagner, 1994; Espinosa-Soto et al., 2004; Enver et al., 2009; Bornholdt and Kauffman, 2019; Raju and Siggia, 2023). From this perspective, the set of all of a GRN’s attractors comprises all the cell types that an organism with that GRN could generate.

Cellular differentiation implies transitions from one stable gene activity pattern to another. We take into account that such transitions are triggered by molecular signals that start a cascade of gene activity changes (Xiong and Ferrell Jr, 2003; Wolpert et al., 2019; Bornholdt and Kauffman, 2019; Barresi and Gilbert, 2023). We thus assessed which attractor-to-attractor transitions a GRN allows: for every one of the GRN’s attractors we perturbed the activity of each gene, one at a time. Starting network dynamics from such perturbed state can eventually lead the system back to the original attractor or to another of the attractors that the GRN admits. We collect this information pertaining to a particular GRN in a different directed graph that we call a *Cell Type Transition Map* (CTTM). Each node in a CTTM represents one of the GRN’s attractors; an arc pointing from one attractor *α* to another attractor *β* indicates that transitions from *α* to *β* are possible via single-gene perturbations (Fig. 1).

**Fig. 1.**
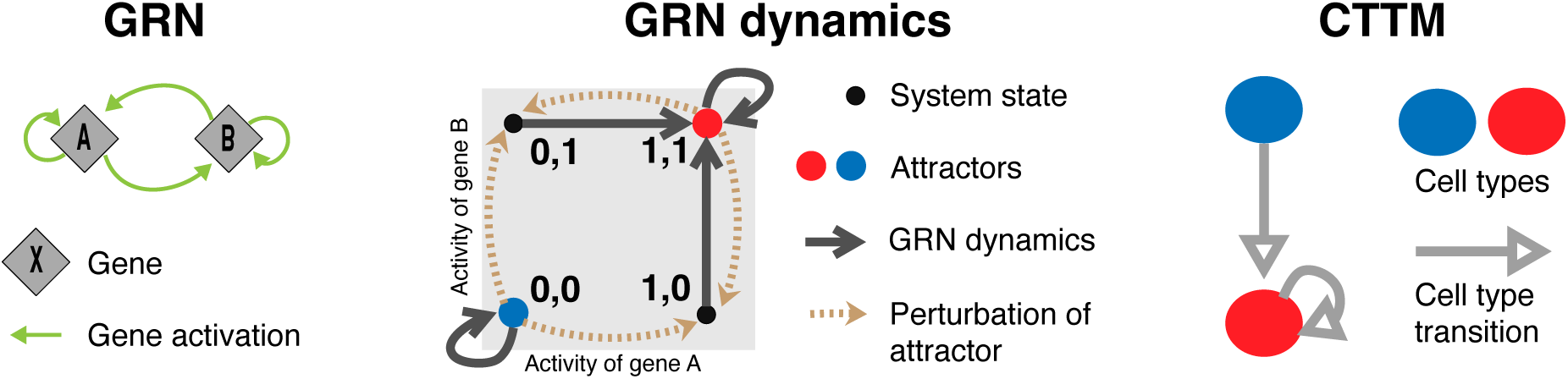
From gene regulatory networks to cell type transition maps. The leftmost panel shows a 2-gene gene regulatory network in which genes A and B self-activate and cross-activate each other. The middle panel shows the corresponding phase space. The four vertices indicate the four possible system states. Gray arrows show which system state follows which, according to the dynamic rules encoded genetically in the GRN. This system has two attractors, marked with a blue and a red dot; the system that reaches any of those states remains there if left unperturbed. Light brown dashed arrows indicate the result of single-gene perturbations of attractors. In this example, all the perturbations of the (1,1) attractor, on red, point to states that eventually lead back to it. In contrast, perturbations of the (0,0) attractor, on blue, take the system to conditions leading to the red attractor. The corresponding Cell Type Transition Map (CTTM) summarizes this information: The GRN allows two cell types and perturbations of the blue cell type lead to the red cell type and perturbations of the red cell type lead back to it.

As our focus is on the simplest regulation patterns that we observe in cell differentiation, we analyzed all the 3*^N^*^2^ possible *N −*gene regulatory networks for *N* values equal to 2, 3 and 4 (Table 1). We only study further those GRNs whose dynamics always lead to stationary equilibrium points; in other words, we consider CTTMs that include one or more oscillatory gene activity patterns as developmental failures and, therefore, non-viable (Wagner, 1996; Siegal and Bergman, 2002; Bergman and Siegal, 2003; Pinho et al., 2014; Le Cunff and Pakdaman, 2014). Still, for the small GRNs that we analyze, most GRNs yield viable CTTMs (Table 1). Many GRNs induce CTTMs that, although viable, are very different to the ones that we expect for the development of multicellular organisms as they contain multiple loops. In fact, many of them consist of a single strongly connected component (Table 1); that is, they form a set of intertwined loops such that, following the direction of the arcs, there is a path from each node to any other node in the set (Harary, 1969; Newman, 2010). If these CTTMs corresponded to a real organism, there would be a differentiation pathway from each cell type to any other, contingent on appropriate sequences of developmental signals; differentiation would be neither progressive nor divergent. Therefore, we do not expect GRNs such as these to play a role in the differentiation of multicellular organisms. Consequently, we focused on those GRNs that yield CTTMs showing the progressivity and hierarchy that characterizes organismal development.

**Table 1.**
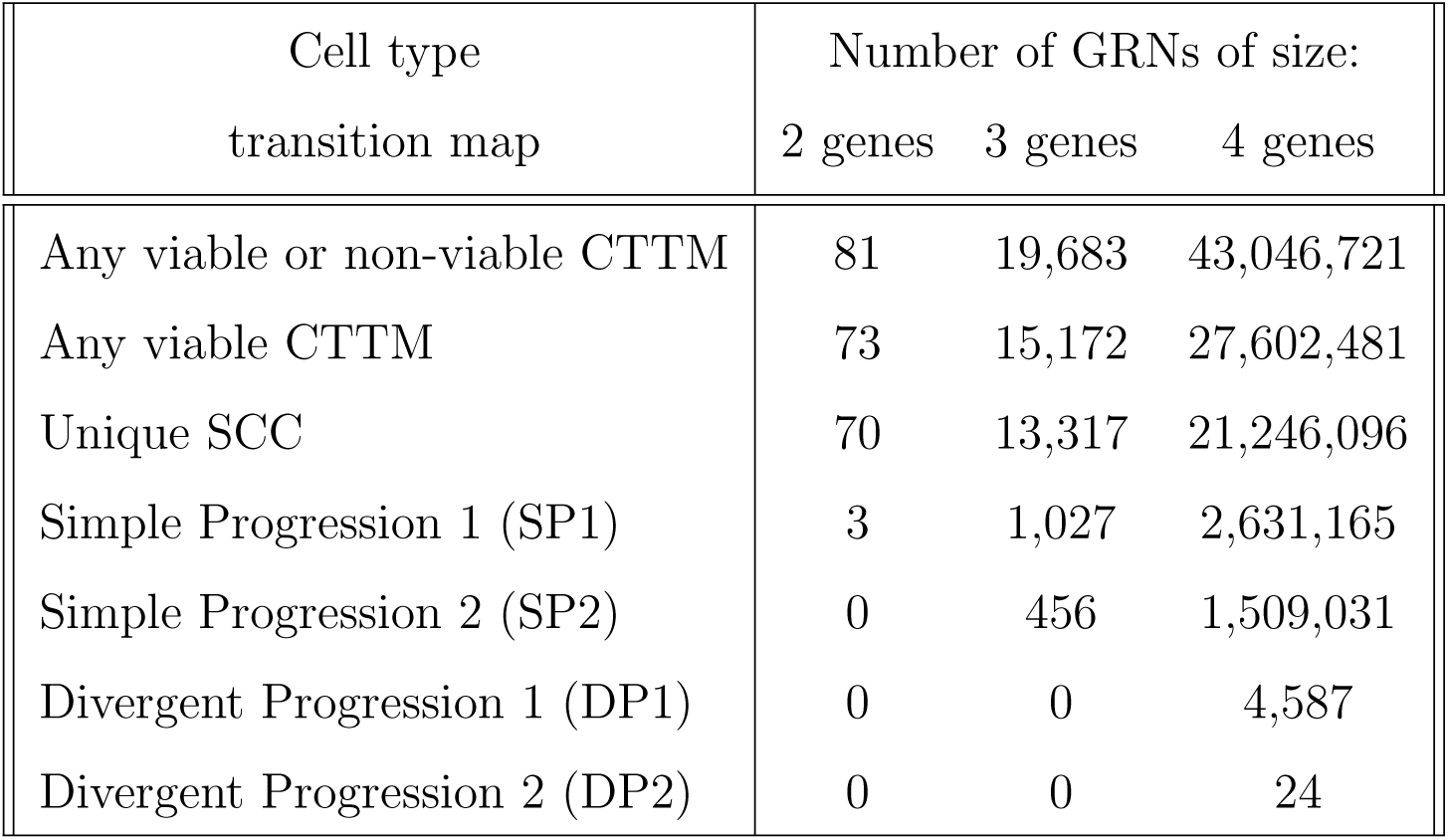
Number of gene regulatory networks that yield specific cell type transition maps.

| Cell type | Number of GRNs of size: |  |  |
|---|---|---|---|
| transition map | 2 genes | 3 genes | 4 genes |
| Any viable or non-viable CTTM | 81 | 19,683 | 43,046,721 |
| Any viable CTTM | 73 | 15,172 | 27,602,481 |
| Unique SCC | 70 | 13,317 | 21,246,096 |
| Simple Progression 1 (SP1) | 3 | 1,027 | 2,631,165 |
| Simple Progression 2 (SP2) | 0 | 456 | 1,509,031 |
| Divergent Progression 1 (DP1) | 0 | 0 | 4,587 |
| Divergent Progression 2 (DP2) | 0 | 0 | 24 |

### Progressive differentiation patterns

First we looked at the simplest progressive transition: a CTTM that contains only two cell types with a transition from one to the other but without the transition in the opposite direction. We call these kinds of CTTMs *Simple Progression* (SP) patterns (Fig. 2). We refer to the node from which the only transition departs in SP CTTMs as the *upstream cell type*. The node that marks the final point of such transition is thus the *downstream cell type*. As there are no transitions away from the downstream cell type, all its perturbations lead back to it; cells that attain such identity become locked-in at it. Some of the properties that we observe on these networks depend on specific details of our GRN models. Still, we can also distinguish traits that do not rely on the model’s idiosincracies. A most important characteristic of GRNs behind SP CTTMs is a positive feedback loop (Fig. 3). The activity state of genes in the loop preserves the identity of the upstream and downstream cell types. This loop thus coordinates all the gene activity changes required to convert an upstream cell type to a downstream cell type. The most resilient state for genes in the loop is associated to the downstream cell type. Consequently, at least some perturbations of loop genes in the upstream cell type lead to the downstream cell type and all the perturbations of the downstream cell type lead back to it, making the transition from upstream to downstream cell types irreversible. We distinguish two classes of SP CTTMs: SP1 CTTMs, in which all the perturbations of the upstream cell type lead to the downstream cell type, and SP2 CTTMs, in which at least one perturbation of the upstream cell type leads back to it (Fig. 2). In our model, only a small fraction of two, three or four-gene GRNs can yield SP CTTMs (Table 1). GRNs that yield SP CTTMs always include a set Γ of genes that have a different activity state in the upstream and downstream cell types. Genes in set Γ are either all active or all inactive in each of the two cell types. At least some of the genes in set Γ are elements of a cycle of positive interactions. Genes in Γ that are not part of such a positive feedback loop are under positive control of genes in the loop. Therefore, this kind of positive feedback loops coordinate the activity of genes in set Γ. Genes outside of set Γ have the same activity state in both the upstream and downstream cell types. Since the system state in which all genes are inactive is always stable in our model, genes outside of set Γ are inactive in both cell types.

**Fig. 2.**
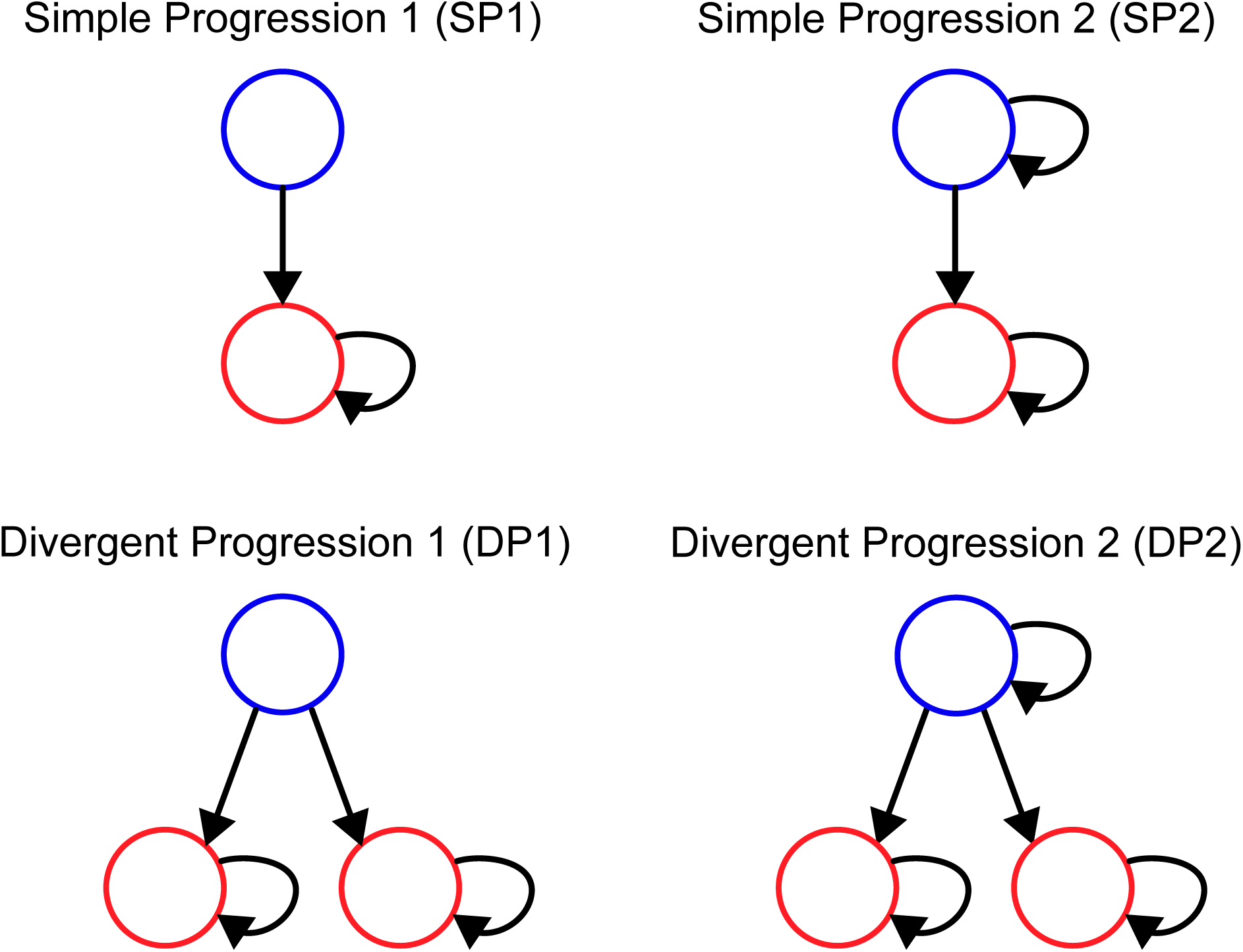
Most common Cell Type Transition Maps (CTTMs) in development. Top panels: Simple Progression (SP) CTTMs. Bottom panels: Divergent Progression (DP) CTTMs. The difference between SP1 and SP2 (or DP1 and DP2) CTTMs is that in the first one, all single-gene perturbations of the upstream cell type lead to a downstream cell type.

**Fig. 3.**
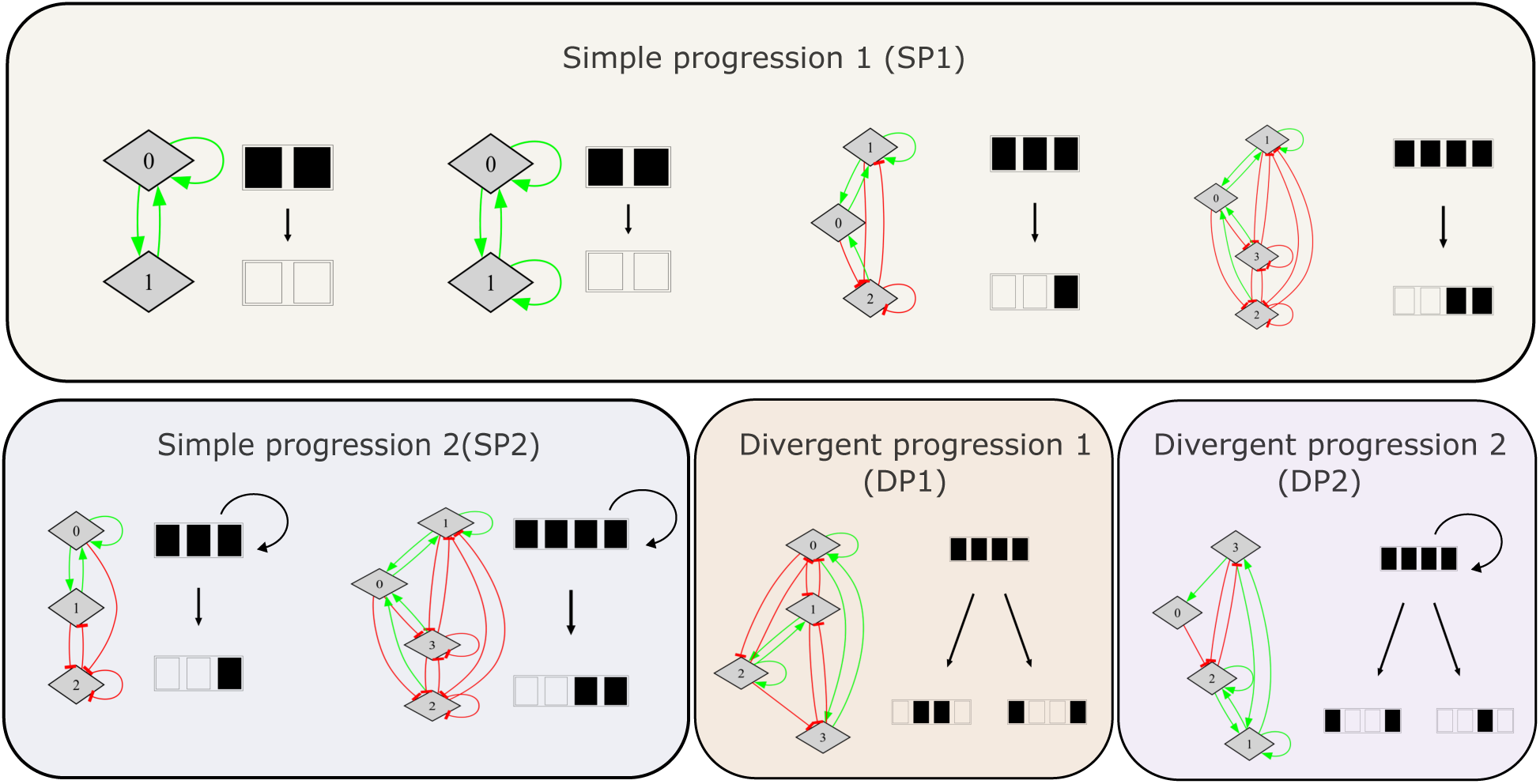
Examples of GRNs that yield simple developmental-like Cell Type Transition Maps. Diamond-shaped nodes represent genes; green arrows and red blunt-end lines indicate, respectively, positive and negative regulatory interactions. The diagram to the right of each GRN represents the CTTM that the GRN yields. In it, each row with *N* squares stands for one of the GRN’s attractors that we associate to a cell type. Whether the *i*-th square in a row is open or black-filled indicates activity or inactivity of the *i*-th gene. Attractors on the top or bottom represent, respectively, the upstream (less differentiated) or the downstream (more differentiated) cell types. Black arrows indicate the cell type transitions that the GRN allows after perturbations of a gene’s activity. With our model, SP1 is the only developmental-like CTTM that two-gene GRNs can produce. To do so, they require cycles of positive interactions also appearing in larger GRNs that yield SP1 and all other developmental-like CTTMs. Larger GRNs that produce SP1 may contain other genes whose transient activation suffices to turn on the genes in the positive cycle of interactions; in these GRNs, any perturbation of the upstream cell type causes a transition to the downstream cell type. GRNs that produce SP2 CTTMs also contain nodes whose perturbation in the upstream cell type bears no consequences. GRNs that yield DP1 or DP2 CTTMs contain two mutually repressing sets of cross-activating genes. The two sets can be non-overlapping or overlapping as in our examples for DP1 and DP2, respectively. In the example for DP2 the two sets overlap in gene 1 which is a differentiation marker that is active in both downstream cell types.

GRNs that produce SP CTTMs in our model can also divide into two classes that we distinguish by the activity of genes in set Γ in the upstream cell type (Supplementary Table S1). In those in which these genes are inactive in the upstream cell type, the cycle of positive interactions must intertwine with at least other cycle of positive interactions of different length (Supplementary Figs. S1,S2). In this case, if all genes in Γ are inactive, as in the upstream cell type, transient activation of some of the genes in the positive loops suffices to eventually turn on all the genes in Γ. That is, genes in Γ produce an all-or-none response to a signal that transiently activates some of them. Moreover, in the downstream cell type, the activity of the whole set Γ is resilient to transient inactivation of anyone of the genes in the set, making the all-or-none response unidirectional.

The other class of model GRNs yielding SP CTTMs are those in which genes in Γ are active in the upstream cell type. In this case, the positive feedback loop is brittle: any transient inactivation of a gene in the loop eventually makes the whole set inactive. Again, this is an all-or-none response. At the same time, transient activation of any gene in the loop eventually wanes if all other genes are initially inactive. Therefore, perturbation of the active loop in the upstream cell type leads to the downstream cell type and perturbation of the inactive loop in the downstream cell type produces no effect: Cell type transitions only occur in one direction.

### Divergent and progressive cell type transition maps

One of the most notable features in development is an undifferentiated cell’s ability to attain any of different alternative cellular fates. Thus, we looked for the simplest *divergent and progressive* (DP) CTTMs. The DP CTTMs contain three different cell types, one is upstream and two are downstream; small perturbations of gene activity in the upstream cell type lead to either of the two downstream cell types. Cell identity becomes locked-in at either of the two downstream cell types: there are no transitions from one to the other or from any of them to the upstream cell type (Fig. 2).

We found that all the GRNs with DP CTTMs contain two distinct sets of genes; genes in either set are active in one of the downstream cell types but not in the other, and therefore one could consider genes in one such set as *cell type marker genes*. Reciprocal inhibitory interactions between genes in the two sets impose such a mutually exclusive activity pattern (Fig. 3).

Stability of cell type marker gene activity in their corresponding downstream cell type requires maintaining their activity through positive feedback. As in SP CTTMs, this arrangement underlies irreversibility of the activation of the genetic program associated to a downstream cell type. Most often, such positive feedback loops involve only genes that are markers of the same downstream cell type. However, in a few GRNs, said feedback loops also implicate one or more genes that are active in the two downstream cell types but inactive in the upstream cell type. We may consider such genes as markers not of a specific cell type but rather as markers of the differentiated state in the two downstream cell types. We refer to them as *differentiation marker genes*; they help to maintain the activity of either mutually exclusive set of genes (Fig. 3).

In some GRNs there may exist genes that are a special kind of cell type marker genes. Like other cell type marker genes, they are active only in one downstream cell type; their peculiarity is that they do not take part in the positive feedback loops that define that cell type. Their activity is due to their positive regulation by other marker genes that do take part in the positive cycle. Because their activity does not affect genes in the cycle, their transient activation by perturbation of a cell type in which they are usually inactive is non-consequential, leading back to the original unperturbed stable activity profile. GRNs that yield DP CTTMs and do not include this kind of futile marker genes produce one specific class of DP CTTMS, DP1, in which any single-gene perturbation of the upstream cell type leads to one of the downstream cell types. In contrast, the existence of these futile marker genes in GRNs that produce DP CTTMs implies another class of such differentiation patterns, DP2, in which there is at least one single-gene perturbation of the upstream cell type that leads back to it (Fig. 2).

We also note some observations that should not be extrapolated to real biological networks as they are peculiarities due to our model’s details. In our model, the smallest GRNs associated to the simplest divergent progressive (DP) differentiation patterns consist of four genes. This condition arises from the model’s need to avoid cyclic attractors, from all interactions having the same weight, and from gene activity being a discrete variable. Hence, maintaining gene activity specific to a downstream cell type requires at least two cycles of positive interactions that converge in one or more of the cell type marker genes, as in GRNs that yield SP CTTMs (Supplementary Fig. S1). Moreover, to avoid the possibility of some perturbations leading to cyclic attractors, the lengths of both such positive feedback loops must be coprime numbers. Such a need for multiple circuits of different length imposes a lower limit on the number of genes that produce DP CTTMs in our model. Moreover, all the GRNs that yield DP CTTMs have all the genes inactive in the upstream cell type. This situation arises because inactivity is the default state of any gene in our model (see Methods), thus reflecting a pervasive condition in the transcription of eukaryotic genes (Struhl, 1999).

### Large sets of GRNs that yield divergent and progressive CTTMs can be traversed through single-mutation steps

Among all possible 3^42^ (about 43 *×* 10^6^) 4-gene GRNs and among the more than 27 *×* 10^6^ of them that produce viable CTTMs, only 4,611 yield a DP CTTM. Specifically, there are only 4,587 and 24 GRNs that produce, respectively, a DP1 or a DP2 differentiation pattern (Table 1). We cannot analyze exhaustively GRNs with more than four genes. In our model, the number of possible GRNs increases with the number of genes *N* as 3*^N^*^2^: there are more than 8 *×* 10^11^ and 1.5 *×* 10^17^ 5- and 6-gene networks, respectively. Thus, we explored the universe of larger networks by random uniform sampling. We found 24 GRNs that yielded DP CTTMs (21 DP1, 3 DP2) in our random sample of 10^5^ 5-gene networks (0.023 *±* 0.01%, 95% confidence level). As for 6-gene GRNs, 35 networks (0.035 *±* 0.01%) in our sample produced DP patterns (26 DP1, 9 DP2). The apparent increase in the fraction of DP-associated GRNs with the number of genes is not indefinite: in a large random sample of 12-gene GRNs we only found 15 DP-producing networks, all of them yielding DP1 CTTMs (0.015 *±* 0.008%).

How can evolution find so often GRNs that underlie DP CTTMs in real organisms when these networks are so scarce? One possibility is that it could be easy to find them when the search starts from other GRNs that already yield DP CTTMs. Indeed, that may be the case. Of the 4,611 gene networks that produce DP CTTMs, 4,491 of them (97.4%) conform three equal-sized sets, each of 1,497 GRNs (see details of configuration of these three large sets and of the remaining 2.6% in Supplementary Tables S2,3). The most important property of any one of these large sets is that it is mutationally connected: the whole set can be traversed through single-mutation steps without ever leaving it. Therefore, an evolving population may explore many genetic configurations without ever losing the ability to yield DP patterns. The distinct GRNs that a population can assay may be very different, allowing it to try very different developmental modes and mechanisms. Indeed, two GRNs sampled at random from one of the large mutationally connected sets differ, on average, by 6.13 interactions; the maximum difference between two networks in the large mutationally connected set equals 12 of 16 possible interactions (Fig. 4A).

**Fig. 4.**
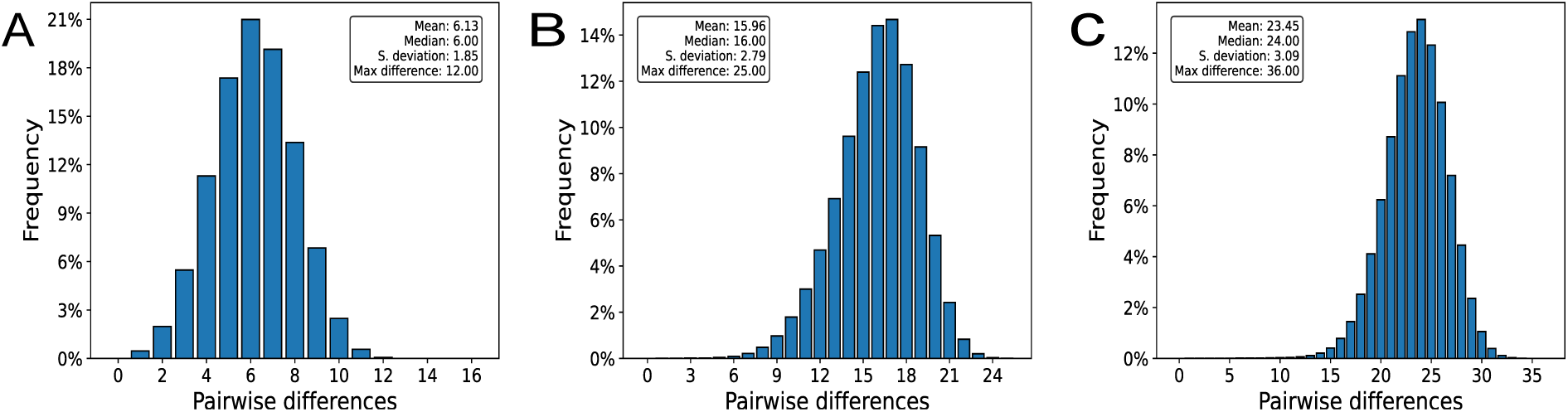
Distribution of pairwise differences between GRNs that yield a DP1 CTTM and that lie in the same mutationally connected set. We calculated the distance between GRNs as the smallest number of changes required to transform one network into another. A. Distribution of pairwise differences between all the 1,497 4-gene GRNs in one of the large mutationally connected sets associated to DP1 CTTMs. B,C. Histograms take into account all the different networks found along a random mutational walk that only allowed mutations that preserved a GRN’s ability to yield a DP1 pattern. The number of mutations along the random walk was 10^5^. B. Five-gene GRNs. C. Six-gene GRNs.

Although we cannot describe the arrangement of all the GRNs that yield DP CTTMs when such networks contain more than four genes, we can ask whether they also tend to comprise large mutationally connected sets. After selecting one random network that yielded a DP1 (or DP2) CTTM, we started a random mutational walk in which we only allowed mutations that preserved the initial DP1 (or DP2) differentiation pattern. To obtain a lower-bound estimate of the size of a mutationally connected set, we counted the number of distinct networks that yielded a DP1 (or DP2) CTTM after 10^5^ mutational steps. For DP1 CTTMs, the number of distinct networks in the random walk was 31,948 and 39,866 for 5- and 6-gene GRNs, respectively. For DP2 CTTMs, the analogous numbers for 5- and 6-gene GRNs were 26,757 and 37,795. The mutationally connected sets span a wide range of network configurations, as shown in Figures 4B,C for DP1 CTTMs and in Supplementary Figure S2 for DP2 CTTMs. The maximum number of interaction differences in the mutationally connected sets for DP1 and for DP2 reach the theoretical maximum of 25 and 36 for 5 and 6 gene networks, respectively: There are pairs of networks in the mutationally connected set that differ in all the possible *N* ^2^ interactions. Together, our results support that gene regulatory networks that produce DP CTTMs are easily navigable through mutation.

### Connection patterns associated with hierarchical CTTMs abound in experimentally sustained developmental GRNs

So far we have described the association of distinct kinds of feedback loops, and their combinations, with hierarchical differentiation patterns. Thus, we asked how frequent were these connection patterns in GRNs as close as we could to those of real multicellular organisms. As explained in Methods, we compiled a database of 55 GRNs involved in cellular differentiation sustained on experimental information and covering a wide range of taxonomic groups.

For each experimentally sustained network in our database, we built 1,000 randomized versions of it using a modification of the configuration model for directed, signed networks (Milo et al., 2003; Newman, 2010). Each of these randomized networks has the same degree distribution as the original network, although interactors are picked randomly and, therefore, usually they are not the same as in the original network. Hence, we exclude changes in the number of connections or a particular degree distribution as a cause for any deviation from randomness in the experimentally sustained networks.

We assessed whether the number of previously identified positive feedback loops patterns was substantially different in the developmental GRNs and in their randomized counterparts. We found that the number of positive feedback loops overall is greater in developmental GRNs than the corresponding average of their respective randomized networks (Fig. 5A). We examined the number of positive feedback loops with a specific length and number of positive interactions. After correction for multiple hypothesis testing using the false discovery rate (Benjamini and Hochberg, 1995), we found that most of the different kinds of positive feedback that we assayed were significantly more abundant in developmental GRNs than as expected by chance (Supplementary Table S4). These cases include direct self-activations, two-gene mutual inhibitions, two-gene reciprocal activations and three-gene cycles that included two repressive interactions. We only found one positive feedback pattern whose frequency was not significantly different from random expectation: three-gene cycles without repressive interactions were, on average, slightly less frequent in developmental GRNs (mean*±*SD: 1.58 *±* 3.26) than in their randomized counterparts (1.92 *±* 2.35). By contrast, we observed that the number of overall negative feedback loops patterns is significantly lower than in the randomized versions (Supplementary Figure S3). The same is true for individual negative feedback loop patterns (Supplementary Table S5).

**Fig. 5.**
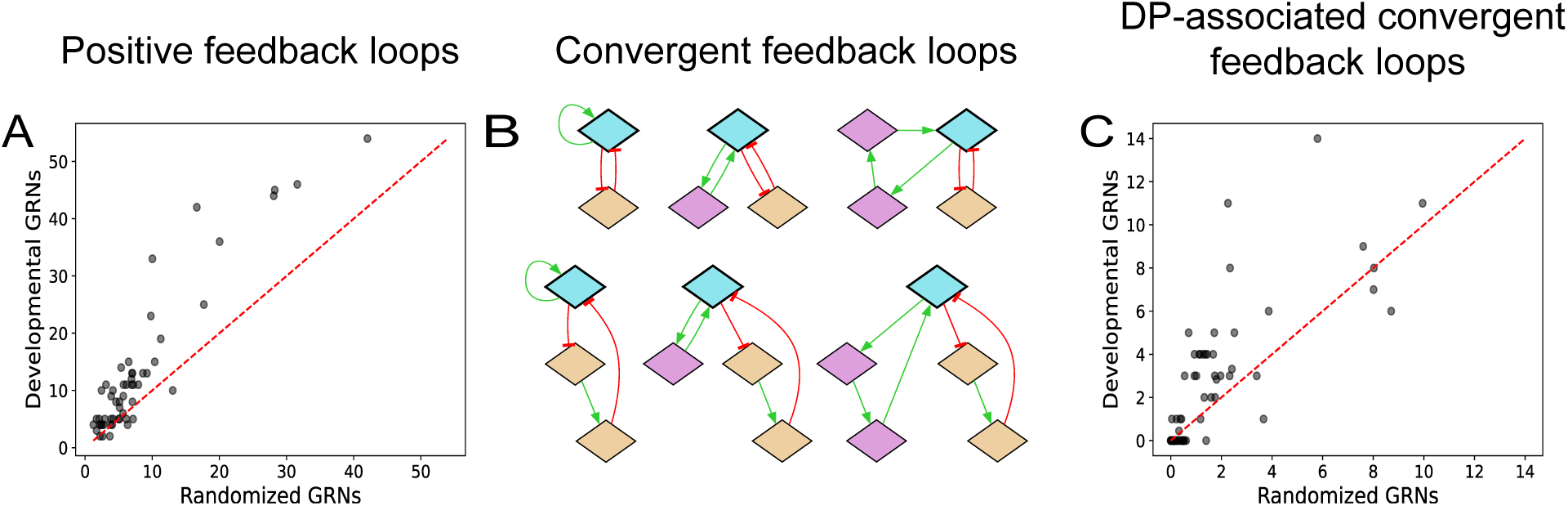
Connection patterns associated with hierarchical CTTMS are more abundant in developmental GRNs than as expected by chance. A. The number of positive feedback loops (including self-activations, reciprocal activations, reciprocal inhibitions and positive 3-gene cycles (Supplementary Table S4) is greater in developmental GRNs than in the average of their randomized counterparts. Dotted diagonal indicates the identity line. Wilcoxon signed-rank test *W* = 1, 429; *p* = 1.7 *×* 10*^−^*^8^. B. Combinations of cycles of positive interactions and mutual inhibition. Positive interactions appear in green and negative interactions in red. The two kinds of positive cycles converge in blue nodes. Cycles of positive interactions traverse pink and blue nodes; mutual inhibition patterns consist of orange and blue nodes. C. Number of genes in which cycles of positive interactions and mutual inhibitions converge (blue nodes in panel B) in developmental GRNs and in their randomized counterparts. *W* = 1, 118; *p* = 0.002.

We have already shown that GRNs that produce divergent and progressive CTTMs always contain specific combinations of positive feedback loops. Namely, DP CTTMs require two distinct kinds of these patterns to converge in one gene: cycles of positive interactions to maintain the expression of genes associated to a downstream cell type, and reciprocal inhibitions with other sets of genes, to exclude activation of genes associated to other downstream cell types. Thus, we assessed whether these combinations appear with a frequency higher than as expected by chance in our collection of developmental GRNs. Limiting ourselves to cycles of length three or shorter, we counted, in each GRN, the number of genes involved in both such kinds of positive cycles (Fig. 5B). We found that developmental GRNs have a greater number of such genes than their randomized counterparts (Fig. 5C). Overall, our results support that the coupled cycles of positive interactions and mutual inhibitions that we identified in GRNs that yield progressive and divergent CTTMs are frequent components of developmental GRNs.

## Discussion

Differentiation pathways usually conform a tree-like pattern in clonal multicellular organisms (Garcia-Bellido and Ripoll, 1978; Kauffman, 1982; Stent, 1985; Zhou and Huang, 2011). This pattern arises because of the scarcity of returns to antecedent states and the frequency of diversifying branching points. How cell type transition maps, CTTMs, acquire such a peculiar organization requires an explanation that must reside in the properties of gene regulatory networks. After all, gene regulatory networks orchestrate changes in gene activity and gene activity patterns define cell types.

A first possibility to explain the commonality of progressive and divergent CTTMs in development would be that hierarchical CTTMs are really a nearly inescapable dynamical feature of gene regulation (Kauffman, 1982). In this scenario, all or the vast majority of GRNs would yield such developmental-like CTTMs. However, this does not seem to be the case. When we looked at all possible 2,3 and 4-gene GRNs we found that most viable GRNs produce a tangle of cycles of cell type transitions. This observation goes along the same lines as previous studies that showed that cycles of cell type transitions are overly abundant in large regular random Boolean networks (Kauffman, 1969; Ribeiro and Kauffman, 2007).

In our exhaustive sample of small GRNs we found a few that yielded *simple progressive* (SP) CTTMs. All the GRNs that had this ability contained a positive feedback loop that guaranteed that cell type transitions only occurred in one direction. This finding supports the role of positive feedback in developmental lock-in (Davidson et al., 2002; Xiong and Ferrell Jr, 2003). Indeed, we did not find any other viable alternative to achieve one-way cell type transitions.

We only found *divergent and progressive* (DP) CTTMs, the most distinctive CTTM of multicellular development, in GRNs with four or more genes. Positive feedback also played an important role in this CTTM: the identity of each of the two downstream, more differentiated, cell types is preserved through positive feedback. This observation may not be surprising since DP patterns are a particular composition of two SP patterns. Moreover, there is mutual repression between the two sets of genes whose activity defines each of the downstream cell types. Hence, the activity of the two sets of genes becomes mutually exclusive. Previous theoretical and empirical work has already defended the importance of self-activating and mutually repressing groups of genes for development (Huang et al., 2007; Enver et al., 2009; Wang et al., 2010; Zhou and Huang, 2011; Bessonnard et al., 2014; Jia et al., 2017; Hari et al., 2022; Shyam et al., 2025; Duddu et al., 2026). For example, Graf and Enver (2009) claim a central role in development for *transcription factor antagonism*, as they call the reciprocal inhibition between sets of cross-activating genes. Our work supports that this kind of arrangement is the only one that allows a small set of genes to yield progressive and divergent cell type transition maps.

Our analysis of model GRNs support that sets of cross-activating genes that reciprocally inhibit other such sets underlie a hierarchical arrangement of cell type transitions. Are our observations pertinent to the organization of differentiation in nature? Ideally, one should assess this question experimentally for a wide diversity of cell types and a broad range of species. Since this is not possible, we settled on what we considered a second best option: experimentally sustained developmental gene regulatory networks available in the scientific literature (see Methods and Supplementary Table S6). We found that positive feedback, be it mutual inhibition or cycles of positive interactions is more abundant in developmental GRNs than expected by chance. In contrast, negative feedback appears less often in such networks. Moreover, in agreement with our findings on model GRNs, genes in which there is convergence of mutual inhibition with some genes and cycles of positive interactions with other genes are also frequent above random expectation in our collection of experimentally sustained developmental networks.

*Character identity networks*, ChINs, are gene regulatory networks that define discrete, individualized, cell (or character) identities and that may be considered evolutionary units (Wagner, 2014; Arendt et al., 2016). Importantly, a ChIN’s genes collaborate to maintain their joint activity that suppress ChINs associated to alternative cellular identities (Wagner, 2014); thus, ChINs conform to the gene regulation patterns behind a progressive and divergent organization of cellular differentiation. The coincidence suggests that hierarchy in cell differentiation and discreteness of cell types may be two sides of the same coin.

Our work shows that only a minute fraction of all possible GRNs can yield developmental-like CTTMs. This may seem like an ineludible obstacle for the evolution of progressive and divergent CTTMs: How can evolution find them? We addressed this question for GRNs that produce DP CTTMs, that best characterize development. We found that the vast majority of DP-yielding GRNs conform large mutationally-connected sets (Wagner, 2008, 2011). In other words, there are long sequences of mutations that can drastically alter a GRN without ever stop producing DP CTTMs. Hence, once evolution finds one such DP CTTM, it is easy to preserve a hierarchical organization of differentiation while still evolving.

Our contribution focused on searching the smallest GRN’s that sufficed to yield the simplest developmental-like CTTMs. Our main goal was the identification of developmental modules whose assembly could lead to the generation of the more complex differentiation patterns that we observe in multicellular organisms. Further modelling work and comparison of its results with available empirical evidence will be useful to understand how these developmental modules collaborate. The same approach may allow us to enquire how the structure of developmental GRNs and the properties of their resulting CTTMs play a role in the evolution of new cell types (Arendt, 2008; Arendt et al., 2016).

## Methods

### Developmental dynamics

To assess how the organization of differentiation emerges from the dynamics of GRNs and their arrangement of regulatory interactions we use a version of a GRN model proposed by Wagner (1994) that has been used to study many aspects of evolution and development associated to GRNs (reviewed in: Fierst and Phillips, 2015; Wang, 2019). The model considers a cell with *N* genes in which the cellular state at time *t* is described by a vector *s^t^ = {s^t^_1_, s^t^_2_, s^t^_3_, …, s^t^_N_*, where s^t^_i_ = 1 if gene *i* is active at time *t*, or s^t^_i_ if gene *i* is inactive at time *t*. A GRN can be represented as a matrix **W** whose entries have three possible values. An entry *w_ij_* = 1 implies that gene *j* promotes the activation of gene *i*; if *w_ij_* = *−*1,then gene *j* represses the activity of gene *i*; otherwise, if *w_ij_* = 0 gene *j* does not affect directly the expression of gene *i*. The activity state of the *N* genes changes over time due to the GRN’s regulatory interactions according to the difference equation:

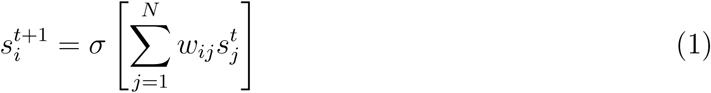

where *σ*(*x*) is a step function that equals 1 when *x >* 0 and equals 0 otherwise. This implies that, in this model, genes without active regulators will be inactive. This behavior matches that of most eukaryotic genes, whose transcription requires the presence of positive regulators (Struhl, 1999).

Starting from an initial condition **s**^0^, iterative application of Eq. 1 leads the system to an attractor that may be a fixed-point or a limit cycle. A fixed-point attractor corresponds to a cell in which all gene activity stops changing, maintaining a steady gene activity pattern. In a limit cycle, gene activity starts fluctuating, following an indefinitely repeating sequence of system states. We associate fixed-point attractors to the stable gene activity patterns that define discrete cell types (Kauffman, 1991; Kauffman et al., 2003; Wagner, 1994; Espinosa-Soto et al., 2004; Enver et al., 2009; Bornholdt and Kauffman, 2019; Raju and Siggia, 2023). As in previous work (Wagner, 1996; Siegal and Bergman, 2002; Bergman and Siegal, 2003; Pinho et al., 2014; Le Cunff and Pakdaman, 2014), we regard lack of stabilization of gene activity as a developmental failure. Hence, we deem GRNs that produce limit cycle as non-viable.

In our model, regulatory interactions coded in the matrix **W** determine all the fixed-point attractors that a GRN can yield and thus, all the cell identities that an organism with such a GRN can produce. Which cell identity a specific cell attains, among all allowable possibilities, depends on the cell’s initial condition **s**^0^. Therefore, we can find a GRN’s repertoire of cell types starting developmental dynamics from each of all possible 2*^N^*initial states.

### Cell type transition maps

We define a Cell Type Transition Map (CTTM) as a directed graph in which nodes represent a GRN’s fixed-point attractors and each arc represents the possibility of a cell type transition in response to perturbation of the activity of a single gene. We assume that perturbation results from molecular signals that reach a cell. In the model, we simulate those perturbations as single-gene activity changes of one of the network’s attractors. To find out which cell type transitions a GRN allows, we first searched for all the network’s attractors by starting dynamics from each of all 2*^N^* possible initial conditions. If the GRN produced one or more limit cycles, the GRN was deemed non-viable and we stopped its analysis. Otherwise, for each attractor *α* we assayed, one at a time, all *N* possible single-gene activity changes. We started network dynamics from each single-gene perturbation of attractor *α*. Dynamics could lead back to the same attractor *α* or to any of the remaining network attractors. If perturbation lead to attractor *β*, we included in the GRN’s CTTM a transition from cell type corresponding to attractor *α* to that corresponding to attractor *β*.

### Mutationally connected sets

We define a mutationally connected set as a set of GRNs that share some property and that can be traversed through single mutation steps without ever leaving the set. Because we analyzed exhaustively all the 4-gene GRNs, we could determine precisely the arrangement and sizes of mutationally connected sets of 4-gene GRNs that yield DP1 or DP2 CTTMs. For larger GRNs we assessed the size of mutationally connected sets associated to DP1 or DP2 CTTMs through approximation. We started a random mutational walk from a network that yielded the desired CTTM, be it DP1 or DP2, and that we had obtained from random uniform sampling. Each step a mutation affected one entry of the GRN’s matrix **W** picked at random uniformly. The new value was one of the entries’ allowable values (−1,0,1) but discarding the possibility of preserving the value that the entry had before mutation. If the new mutant network still produced the initial CTTM, we accepted the mutation and logged the new GRN. Otherwise, the mutation was rejected. The size of the GRN log, after excluding repeated instances, was a lower bound estimate of the CTTM’s mutationally connected set.

### Assembling a database of developmental GRN models

We gathered experimentally and computationally supported GRN models involved in cell fate determination in animals and plants. We compiled such a list from the model databases The Cell Collective (Helikar et al., 2012), Biodivine Boolean Models (Pastva et al., 2023), GINsim (Chaouiya et al., 2012) and Kadelka et al.’s study (2024), as well as Peter and Davidson’s book (2015) and works that we had previously read (see Supplementary Table S6). Whenever multiple works addressed the same biological system, we only considered the most complete for our analysis. Our collection comprises 55 developmental GRN models available to us as of October 2025.

### Generation of randomized networks for each GRN model

For each gene regulatory network in our database, we built 1,000 randomized versions of it using a modification of the configuration model for directed, signed networks (Newman, 2010; Milo et al., 2003). Specifically, for each randomized network, we first log the number of incoming and outgoing interactions for each node in the original network. Then, after erasing all the network’s interactions, we iteratively propose random interactions accepting them only if the prospect target (source) node does not already surpass its pre-defined number of incoming (outgoing) interactions. Finally, we pick some interactions at random and turn them into negative interactions, to also preserve the number of positive and negative interactions of the original network. Each of the resulting randomized networks has the same number of nodes, positive and negative interactions, and the same degree distribution as the original network.

The code required to replicate our analyses is available at GitHub: https://github.com/cespinosas/Differentiation2026.

## Acknowledgements

We thank J.C. Sánchez-Leaños for technical assistance. C.E.-S. acknowledges financial support from SECIHTI Mexico (CBF-2025-I-98). L.F.M.-S. (824130) and A.Ch.G.-V. (4001820) received student scholarships from CONAHCyT, Mexico.

