## Supplementary Figures for "Gene regulatory networks and the organization of cell differentiation in multicellular organisms"

L. Fernando Meraz-Segura[1], A. Chárbel Guerrero-Vaca[1], Rafael Alcaraz Barrera[1], and Carlos Espinosa-Soto*[1]

[1]Instituto de Física, Universidad Autónoma de San Luis Potosí, Av. Parque Chapultepec 1570, San Luis Potosí, Mexico

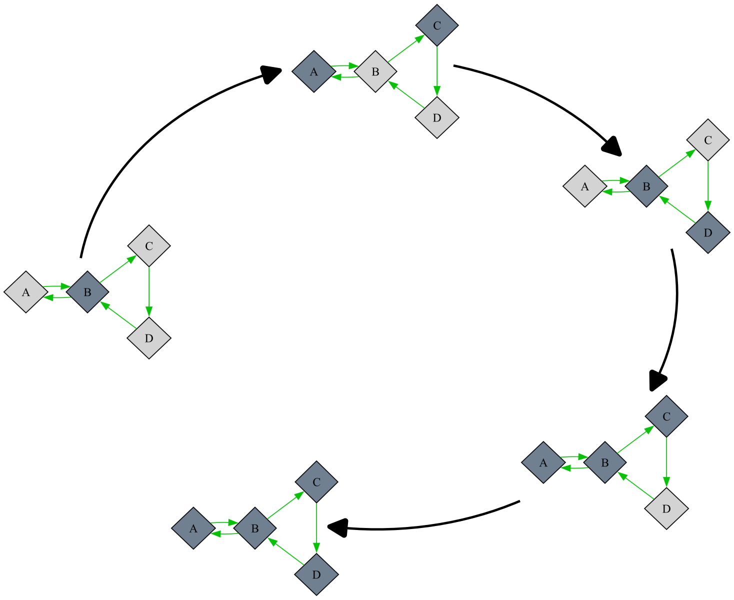


**Fig. S1. Maintaining the activity of a set of genes after transient perturbation of one of them requires the convergence of two cycles of positive interactions.** The length of the two cycles must be coprime numbers; otherwise, transient activation of a single gene leads, in our model, to ever-repeating oscillations of gene activity. Active genes appear in dark gray. In the initial condition only gene $B$ is active; one might attain such an initial condition from a single-gene perturbation of the attractor in which all genes are inactive.

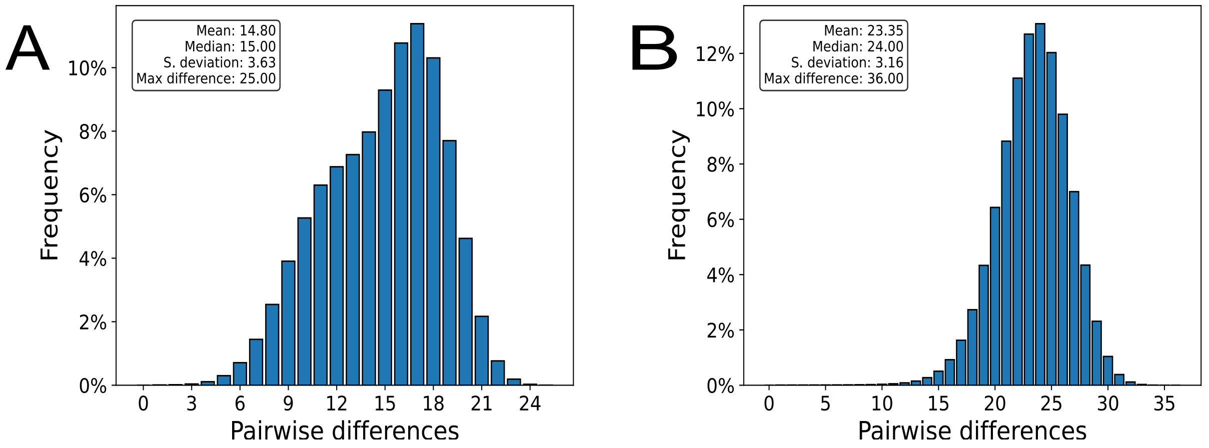


**Fig. S2. Distribution of pairwise differences between GRNs that yield a DP2 CTTM and that lie in the same mutationally connected set.** We calculated the distance between GRNs as the smallest number of changes required to transform one network into another. The sample includes all the different networks found along a random mutational walk that only allowed mutations that preserved a GRN's ability to yield a DP2 pattern. The number of mutations along the random walk was $10^5$. A. Five-gene GRNs. B. Six-gene GRNs.

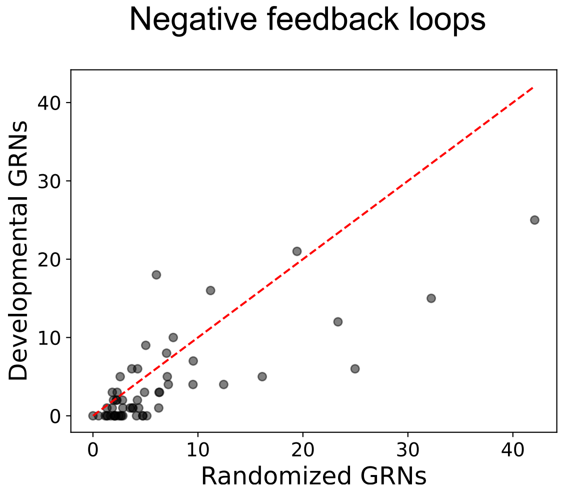


**Fig. S3. The number of negative feedback loops is significantly lower in developmental GRNs than in their randomized counterparts.** Dotted diagonal indicates the identity line. Wilcoxon signed rank test. $W = 269$; $p = 2.3 \times 10^{-5}$.
