## Supplementary Tables for "Gene regulatory networks and the organization of cell differentiation in multicellular organisms"

L. Fernando Meraz-Segura[1], A. Chárbel Guerrero-Vaca[1], Rafael Alcaraz Barrera[1], and Carlos Espinosa-Soto*[1]

[1]Instituto de Física, Universidad Autónoma de San Luis Potosí, Av. Parque Chapultepec 1570, San Luis Potosí, Mexico

**Table S1. Description and number of GRNs that yield SP CTTMs.** The default attractor is the equilibrium point in which all genes are inactive.

| Position of default attractor | Description | CTTM | Number of GRNs with: 3 genes | 4 genes |
|---|---|---|---|---|
| Upstream | Activation of any gene eventually activates the positive feedback loop that sustains gene activity in the downstream cell type. | SP1 | 961 | 2,430,117 |
| | | SP2 | 456 | 1,438,362 |
| Downstream | Conditions leading to the upstream cell type require the activity of multiple genes in the positive feedback loop within set $A$. | SP1 | 66 | 201,048 |
| | | SP2 | 0 | 70,668 |

**Table S2. Arrangement of DP1-associated GRNs in mutationally connected sets.** A mutationally connected set is a maximal set of GRNs with some common property and that can be traversed through single mutation steps without ever leaving the set. 4,491 GRNs belong to one of 3 equal-sized mutationally connected sets. These 3 large sets are isomorphic: there is a one-to-one relationship between GRNs in one set and GRNs in another. Besides yielding DP1 CTTMs, GRNs in one such large mutationally connected set have something in common: Each of their genes never joins two other specific genes as marker genes of the same downstream cell type. Thus, in one mutationally connected set, for example, gene 1 is either a marker gene for a downstream cell type jointly with gene 2, or it is the only marker for a downstream cell type or it is a general differentiation marker active in both downstream cell types. In this set, gene 1 would never be a marker gene for a downstream cell type jointly with neither gene 3 nor gene 4.

| Label of mutationally connected set | Number of GRNs in each mutationally connected set | Number of such mutationally connected sets | Total number of GRNs |
|---|---|---|---|
| A | 1,497 | 3 | 4,491 |
| B | 3 | 24 | 72 |
| C | 1 | 24 | 24 |
| | | Total: | 4,587 |

**Table S3. Arrangement of DP2-associated GRNs in mutationally connected sets.** There are only 24 4-node GRNs that yield DP2 differentiation patterns. These 24 GRNs are not connected to each other; however each of them is connected via a single mutation to one of the small mutationally DP1 connected sets that consist of a single GRN (last row in Table S2).

| Label of mutationally connected set | Number of GRNs in each mutationally connected set | Number of such mutationally connected sets | Total number of GRNs |
|---|---|---|---|
| D | 1 | 24 | 24 |
| | | Total: | 24 |

**Table S4. Number of positive feedback loops in developmental GRNs and in their randomized counterparts.**

| Interaction pattern | Models | | Randomized | | One-tailed Wilcoxon test. $W$ (p-value) | Corrected $p$ (FDR) |
|---|---|---|---|---|---|---|
| | Mean | SD | Mean | SD | | |
| Self-activation | 3.13 | 2.63 | 1.73 | 0.64 | 1,201 ($4 \times 10^{-5}$) | $1 \times 10^{-4}$ |
| Reciprocal inhibition | 2.38 | 2.83 | 0.64 | 0.93 | 1,314 ($4.4 \times 10^{-7}$) | $2.2 \times 10^{-6}$ |
| Reciprocal activation | 2.44 | 2.5 | 1.65 | 1.26 | 1,057 ($3.4 \times 10^{-3}$) | $5.7 \times 10^{-3}$ |
| 3-gene cycle (2 negative interactions) | 3.38 | 6.26 | 2.1 | 4.06 | 984 ($8.8 \times 10^{-3}$) | 0.01 |
| 3-gene cycle (all positive interactions) | 1.58 | 3.26 | 1.92 | 2.35 | 443 (0.99) | 0.99 |

**Table S5. Number of negative feedback loops in developmental GRNs and in their randomized counterparts.**

| Interaction pattern | Models | | Randomized | | One-tailed Wilcoxon test. $W$ (p-value) | Corrected $p$ (FDR) |
|---|---|---|---|---|---|---|
| | Mean | SD | Mean | SD | | |
| Self-inhibition | 0.4 | 1.68 | 0.97 | 0.64 | 140 ($1.1 \times 10^{-7}$) | $4.4 \times 10^{-7}$ |
| Activation-inhibition | 0.85 | 1.4 | 1.78 | 1.79 | 235 ($1.07 \times 10^{-5}$) | $1.43 \times 10^{-5}$ |
| 3-gene cycle (One negative interaction) | 1.73 | 2.92 | 3.15 | 4.66 | 163 ($5.1 \times 10^{-7}$) | $1.03 \times 10^{-6}$ |
| 3-gene cycle (All negative interactions) | 0.96 | 2.33 | 0.595 | 1.35 | 487 (0.02) | 0.02 |

### Table S6. List of developmental GRN models analyzed

| GRN model | Source | Taxon |
|---|---|---|
| Flower development | [12] | *Arabidopsis thaliana* |
| Heart field identity | [22] | *Mus musculus* |
| B cells differentiation | [34] | Mammals |
| CD4+ T cells differentiation | [29] | Mammals |
| Cortical area development | [17] | *Mus musculus* |
| Monocytes differentiation | [37] | *Homo sapiens* |
| Lymphopoiesis | [32] | Mammals |
| Mesoderm specification | [31] | *Drosophila melanogaster* |
| Lymphoid-myeloid specification | [10] | Mammals |
| Gonadal sex determination | [42] | *Homo sapiens* |
| T lymphocytes differentiation | [30] | Mammals |
| Vulva differentiation | [51] | *Caenorhabditis elegans* |
| Eggshell patterning | [13] | *Drosophila melanogaster* |
| Gap genes system | [43] | *Drosophila melanogaster* |
| Pair-rule genes system | [44] | *Drosophila melanogaster* |
| Abaxial-adaxial determination | [54] | *Arabidopsis thaliana* |
| Angiogenesis | [50] | Mammals |
| Cardiomyocytes differentiation | [3] | *Mus musculus* |
| Primary sex determination | [47] | *Gallus domesticus* |
| Dorsal-ventral determination in the imaginal disc | [18] | *Drosophila melanogaster* |
| Primary sex determination | [46] | Eutherians |
| Ectoderm specification | [15] | *Paracentrotus lividus* |
| Endomesoderm specification | [39] | *Strongylocentrotus purpuratus* |
| GMP cells differentiation | [41] | Mammals |
| Embryonic stem cells fate | [36] | *Homo sapiens* |
| Pluripotent stem cells fate | [52] | *Mus musculus* |
| Adipogenesis | [45] | Mammals |
| Root epidermis differentiation | [7] | *Arabidopsis thaliana* |
| Root stem cell niche fate | [16] | *Arabidopsis thaliana* |
| Cardiac progenitor cells fate | [53] | *Homo sapiens* |
| Cardiac cell-fate | [19] | *Xenopus* sp. |
| Chondrocyte differentiation | [24] | *Mus musculus* |
| Early development cell-fate | [8] | *Mus musculus* |
| Sepal primordium polarity | [26] | *Arabidopsis thaliana* |
| Hematopoiesis | [21] | Mammals |
| Inner cell mass cells differentiation | [5] | *Mus musculus* |
| Mesenchymal differentiation | [23] | *Homo sapiens* |
| Keratinocytes differentiation | [11] | *Homo sapiens* |
| Dehiscence zone formation | [33] | *Arabidopsis thaliana* |
| Myeloid progenitors differentiation | [25] | Mammals |
| Natural Killer cells differentiation | [27] | Mammals |
| Pancreas cell fate | [55] | *Mus musculus* |
| Segment polarity genes system | [2] | *Drosophila melanogaster* |
| Ventral spinal cord development | [28] | Vertebrates |
| Th cell types determination | [35] | Vertebrates |
| Vascular bundles cell-fate | [4] | *Arabidopsis thaliana* |
| Mesoderm specification | [6] | *Xenopus laevis* |
| Mesoderm specification | [6] | *Ambystoma mexicanum* |
| Neural patterning | [49] | *Mus musculus* |
| Limb morphogenesis | [40] | Vertebrates |
| Eyes patterning and specification | [1] | *Drosophila melanogaster* |
| Hindbrain segmentation | [48] | Vertebrates |
| Pre-placodal territory and neural crest determination | [20] | Vertebrates |
| Lens development | [38] | Vertebrates |
| Apical neurogenesis | [14] | Bilateria |
| Trophectoderm and endoderm lineage determination | [9] | *Mus musculus* |
